# Zinc Metabolic Defect of Aging Alveolar Progenitors in Progressive Pulmonary Fibrosis

**DOI:** 10.1101/2020.07.30.229567

**Authors:** Jiurong Liang, Guanling Huang, Xue Liu, Forough Taghavifar, Ningshan Liu, Yizhou Wang, Nan Deng, Changfu Yao, Ankita Burman, Ting Xie, Simon Rowan, S. Samuel Weigt, John Belperio, Barry Stripp, William C. Parks, Dianhua Jiang, Paul W. Noble

**Affiliations:** Department of Medicine and Women’s Guild Lung Institute, Cedars-Sinai Medical Center, Los Angeles, CA, USA; Genomics Core, Cedars-Sinai Medical Center, Los Angeles, CA, USA; Department of Medicine, University of California at Los Angees (UCLA), Los Angeles, CA, USA; Department of Biomedical Sciences, Cedars-Sinai Medical Center, Los Angeles, CA, USA

**Keywords:** IPF, aging, zinc metabolism, progenitor, fibrosis

## Abstract

Idiopathic pulmonary fibrosis (IPF) is a fatal form of interstitial lung disease and aging has been identified as a risk factor to the disease. Alveolar type II cells (AEC2s) function as progenitor cells in the lung. Growing evidences indicate that IPF results from repeating AEC2 injury and inadequate epithelial repair. We previously reported that there was a significant loss of alveolar progenitors in the lungs of patients with IPF. In our current study, we performed single cell RNA-seq of epithelial cells from lungs of patients with IPF and healthy donors as well as epithelial cells from old and young mouse lungs with bleomycin injury. We identified a defect of zinc metabolism of AEC2s from IPF lungs and bleomycin-injured old mouse lungs. We further discovered that a specific zinc transporter ZIP8 was down regulated in IPF AEC2s and AEC2s from aged mice. Loss of ZIP8 expression is associated with impaired AEC2 renewal through sirtuin signaling in aging and IPF. Targeted deletion of Zip8 in murine AEC2 compartment led to reduced AEC2 renewal capacity, impaired AEC2 recovery, and worsened lung fibrosis after bleomycin injury. In summary, we have identified novel metabolic defects of AEC2s during aging and in IPF which contribute to the pathogenesis of lung fibrosis. Therapeutic strategies to restore critical components of these metabolic programs could improve AEC2 progenitor activity and mitigate ongoing fibrogenesis.

**In Brief:** Liang et al. performed single cell RNA-seq (scRNA-seq) of epithelial cells in IPF and in mice and discovered a zinc metabolic defect of alveolar progenitor cells (AEC2) in IPF and injured old mice characterized by down-regulation of a specific zinc transporter ZIP8. Manipulation of ZIP8 and zinc in 3D organoid culture of AEC2 in vitro and targeted deletion of Zip8 in AEC2s in vivo demonsrated a role of ZIP8 in promoting AEC2 progenitor function.

**Highlights:**

- ScRNA-seq revealed dysregulation of zinc metabolism in AEC2s and decreased stem cell signaling in IPF
- Reduced *SLC39A8* and related gene expression of IPF AEC2s and aged mouse AEC2s
- ZIP8-dependent zinc metabolism is required for AEC2 renewal
- Targeted deletion of *Slc39a8* impaired AEC2 renewal and promoted lung fibrosis

## INTRDUCTION

Despite extensive efforts, the mechanisms controlling progressive tissue fibrosis remain poorly understood, resulting in the unfortunate lack of effective therapies for patients with progressive tissue fibrosis. Idiopathic pulmonary fibrosis (IPF) is a fatal form of interstitial lung disease. The hallmark of pathogenesis of IPF is repeated epithelial cell injury and inadequate alveolar epithelial repair that leads to excessive fibroblast activation and lung fibrosis (Jiang et al., 2020; Noble et al., 2012).

Type 2 alveolar epithelial cells (AEC2s) function as progenitor cells that maintain epithelium homeostasis and repair damaged epithelium after injury (Barkauskas et al., 2013; Desai et al., 2014; Hogan et al., 2014). We have showed previously that AEC2s were reduced in the lungs of patients with IPF. The remaining AEC2s in the IPF lung have impaired progenitor function and failed to renew themselves adequaly (Liang et al., 2016). However, the molecular mechanisms that control AEC2 stem cell renewal and unremitting lung fibrosis remain poorly understood.

Aging is one of the critical risk factors for IPF. The incidence, prevalence, and mortality of IPF all increase with aging (Raghu et al., 2016; Selman and Pardo, 2014) (Rojas et al., 2015; Thannickal, 2013). Aging has also been linked to lung fibrosis in animal models (Bueno et al., 2018). Phenotypes of cellular aging including epithelial cell apoptosis (Korfei et al., 2008; Lawson et al., 2011), autophagy (Larson-Casey et al., 2016; Sosulski et al., 2015), senescence (Minagawa et al., 2011), telomere shortening (Alder et al., 2008; Naikawadi et al., 2016), mitochondria dysfunction (Bueno et al., 2015; Ryu et al., 2017), and oxidative stress (Anathy et al., 2018) are observed in IPF lungs. Although the concept that IPF is a disease of aging has been well accepted (Martinez et al., 2017; Selman et al., 2016; Thannickal, 2013), there are limited studies focusing on the age-related mechanisms that contributes to the development and progression of IPF. In this study, we dissected the detrimental role of aging in AEC2 progenitor renewal in IPF.

Multiple metabolic dysfunctions undergo with aging (Bueno et al., 2018; Shyh-Chang and Ng, 2017; Tevy et al., 2013; Yaku et al., 2018). Recent studies have linked metabolic changes with stem cell fate (Lee and Hong, 2020; Shyh-Chang and Ng, 2017; Zhang et al., 2018). Zinc is an essential element and involvels in maintaining multiple functions of human body (Bogden, 2004). In the lung, reports showed that zinc deficiency exacerbated ventilation-induced lung injury (Boudreault et al., 2017; Chen et al., 2012b). Patients with acute respiratory distress syndrome (ARDS) showed significant lower plasma zinc levels relative to control individuals (Boudreault et al., 2017; Cander et al., 2011). Zinc deficiency increased lung and spleen damage (Knoell et al., 2009) in animal sepsis models. These evidence suggests the importance of zinc metabolism in lung biology. Zinc homeostasis is controlled by two zinc transporter families, Zn transporters (ZnT) and Zrt-, Irt-related proteins (ZIP) in maintaining zinc mobilization and compartmentalization across biological membranes (Kambe et al., 2015). Studies showed that zinc and zinc transporters played an important role in maintenance homeostasis and progenitor function of intestinal epithelium (Ohashi et al., 2019). Zinc is required for sirtuin signaling which is a major cell energy metabolism pathway (Imai and Guarente, 2014; Yaku et al., 2018). Studies showed that sirtuins play a role in stem cells maintenance and differentiation (Nogueiras et al., 2012; O’Callaghan and Vassilopoulos, 2017). The role of zinc metabolism in lung stem cell renewal and epithelial repair is largely unknown.

In the current study, using unbiased single cell RNA-sequencing (scRNA-seq), we have found dysregulated metabolic pathways of AEC2s from patients with IPF. We discovered a significant decrease of a zinc transporter *SLC39A8* (ZIP8) in IPF AEC2s. We further demonstrated that loss of ZIP8 in IPF AEC2s resulted in impaired progenitor function of the cells via SIRT1. Exogenous zinc treatment increased ZIP8 and SIRT1 expression and promoted renewal capacity of AEC2s. With mouse model of aging, we have found that AEC2s from bleomycin injured aged mouse lungs shared the similar genomic changes and cellular phenotypes with IPF AEC2s. AEC2s from the lungs of aged mice showed reduced Zip8 expression and decreased renewal capacity relative to that of AEC2s from the lungs of young mice. Exogenous zinc was able to promote renewal and differentiation of murine AEC2s while the response to zinc treatment blunted with aging. With genomic modified mice, we further demonstrated that targeted deletion of ZIP8 in AEC2s compartment impaired AEC2s recovery and worsened lung fibrosis after bleomycin injury. To our knowledge this is the first evidence to show that zinc transporter ZIP8 and zinc metabolism play a role in regulating AEC2 progenitor renewal and lung fibrosis.

## RESULTS

### ScRNA-seq revealed the loss of AEC2s in IPF lungs

In the previous study, we found a significant reduction of AEC2 population in patients with IPF (Liang et al., 2016). The remaining AEC2s in the diseased lung had impaired renewal capacity compared to that of the AEC2s from healthy donor lung (Liang et al., 2016). To further investigate the genomic changes of AEC2s in IPF, we performed scRNA-seq of flow cytometry-enriched epithelial (Lin–EpCAM_+_) cells from lung tissues of IPF patients and healthy donors.

We processed single cell homogenates of lungs from six individuals in each group and that contained 11,381 cells from healthy individuals and 14,687 cells from IPF patients, after removing cells with low quality of RNA. The Uniform Manifold Approximation and Projection (UMAP) plots were generated with a Seurat package and the cells from all samples were overlapped very well (Figure S1A, S1B). The major lung epithelial cell types, AEC2, AEC1, basal cell, Club cell, ciliated cell, pulmonary neuroendocrine cell (PNEC), and proliferative (cycling) cells were readily identified with classical cell markers (Figure S1C).

When we compared the cells from IPF lungs to the cells form healthy lungs, we observed much fewer cells in AEC2 cluster of IPF lungs compared to that of the healthy lungs (Figure 1A, 1B). Increased basal-like cells and ciliated cells were observed with the cells from IPF lungs (Figure 1A, 1B), consistent with recent reports (Adams et al., 2020; Habermann et al., 2020; Xu et al., 2016). The investigation into the biological significance of the accumulation of basal and ciliated cells in IPF pathogenesis is under way in the laboratory. In this study, we focus on the investigation of molecular mechanism which causes dysfunction of AEC2 progenitors in IPF. Expression of AEC2 marker genes including *SFTPC, SFTPA2, SFTPB*, and *ABCA3* were all down regulated in IPF AEC2s relative to healthy AEC2s (Figure 1C). We confirmed decreased *SFTPC* expression of flow sorted AEC2s from IPF lungs relative to AEC2s from healthy donor lungs using real time qPCR (Figure 1D).

**Figure 1.**
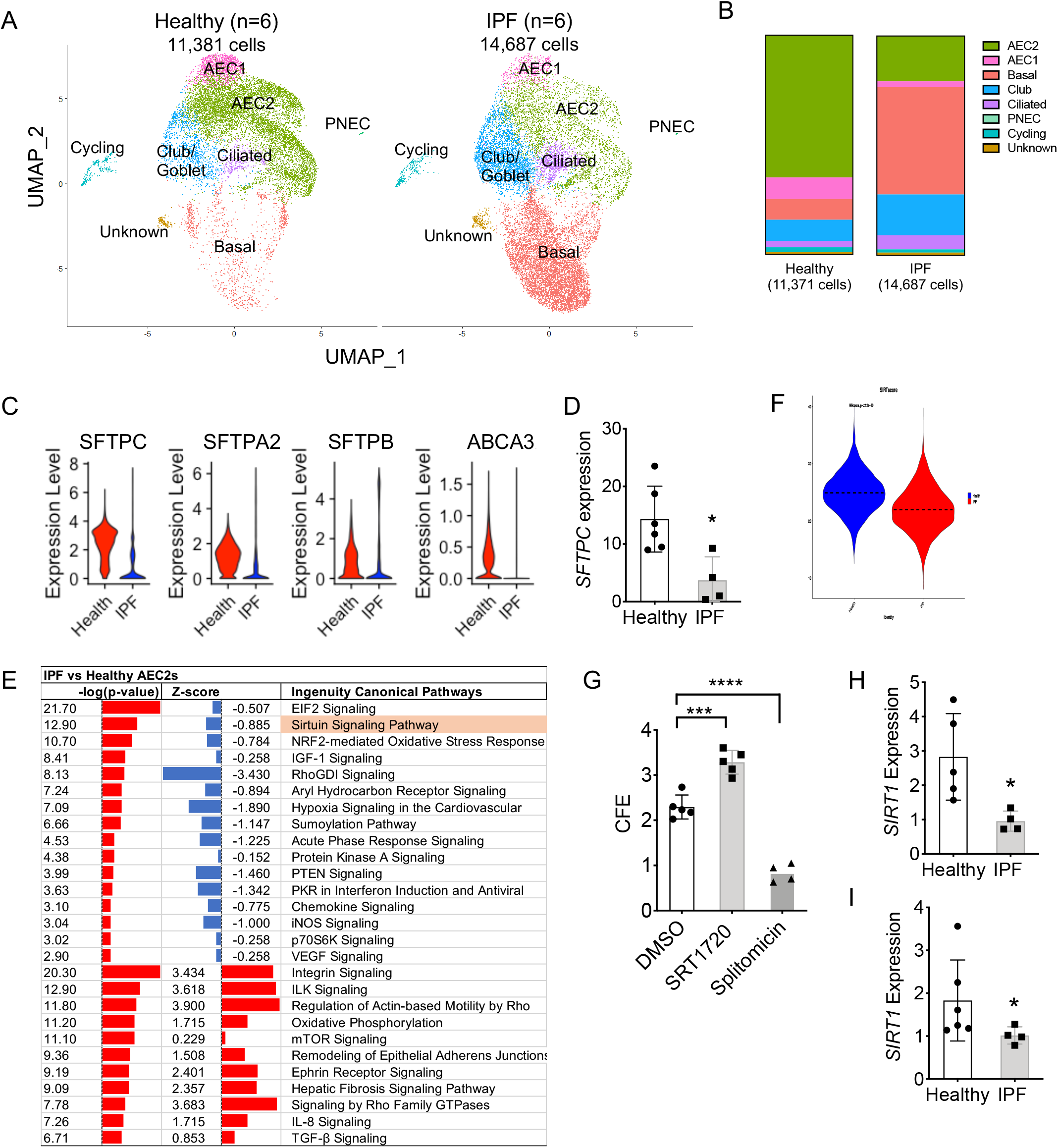
scRNA-seq identifies dysregulation of metabolism and stem cell signaling in IPF AEC2s. (A) UMAP plots of flow enriched EpCAM_+_CD31_−_CD45_−_ cells from healthy (11,381 cells, n = 6) and IPF lungs (14,687 cells, n = 6). (B) Distribution of epithelial cell types. (C) Expression of AEC2 marker genes from healthy and IPF lungs. (D) *SFTPC* expression in healthy and IPF AEC2s by RT-PCR (n = 6 and 4, *p < 0.05). (E) IPA pathway analysis of AEC2s from healthy and IPF lungs. (F) Sirtuin activation score of healthy and IPF AEC2s. (G) 3D organoid culture of AEC2s from healthy lungs treated with SRT1720 and splitomicin (n = 4 – 5, ***p < 0.001, ****p < 0.0001). (H,I) *SIRT1* expression in fresh isolated AEC2s (H) and AEC2s derived from 3D organoids (I) by qPCR (n = 4 - 6, * p < 0.05).

### Dysregulation of metabolism and stem cell signaling in IPF AEC2s

Next, we performed the pathway analysis to compare gene expression of AEC2s from IPF lungs and healthy lungs and found that several metabolic pathways were dysregulated. Based on the findings of our previous study that the AEC2 progenitor function faltered in IPF lungs, we would expect to see the dysregulated stem cell/progenitor cell related signaling in IPF AEC2s. Indeed, we found that the sirtuin signaling pathway was on the top of the list of down regulated pathways of IPF AEC2s (Figure 1E). IPF AEC2s showed lower SIRT score (Figure 1F). The sirtuin signaling is considered as metabolic master switches that control several aspects of metabolism, including glycolysis and fatty acid, lipid, and redox processes (Hershberger et al., 2017). Pathways of NRF2-mediated oxidative stress response and glutathione redox reactions I were down regulated in IPF AEC2s. Down regulation of PTEN signaling in IPF AEC2s (Figure 1E) was consitent with the previous reports (Tian et al., 2019).

The function of the sirtuin signaling has been linked to stem cell renewal and differentiation (Nogueiras et al., 2012; O’Callaghan and Vassilopoulos, 2017). To test the role of sirtuin signaling in regulating AEC2 progenitor renewal, we applied Sirtuin activator, SRT1720, and sirtuin inhibitor, splitomicin, to 3D organoid culture of AEC2s. Our results showed that SRT1720 promoted and splitomicin inhibited colony formation of both human (Figure 1G) and mouse AEC2s (Figure S2A,2B), suggesting sirtuin signaling is important for AEC2 progenitor renewal. Sirt1 is the most important member of sirtuin family (Leibiger and Berggren, 2006; Price et al., 2012). We found decreased expression of SIRT1 in IPF AEC2s both fresh isolated from lung tissues (Figure 1H) and derived from 3D cultured organoids (Figure 1I) relative to that of the AEC2s from healthy lungs using qPCR. These data suggest that dysregulated metabolic programs of IPF AEC2s leads to progenitor function failure of the cells, and lower SIRT activity in IPF AEC2s may contribute to their lower renewal potential.

### Reduced expression of *SLC39A8* and zinc metabolism related genes of IPF AEC2s

Sirtuin signaling is zinc dependent cell energy metabolism pathway, and SIRT enzymatic activities require an adequate level of intracellular zinc (Nogueiras et al., 2012). Next we looked at the expression of zinc metabolism related genes of AEC2s from IPF patient and healthy donor lungs in our single cell RNA-sq data set. We found several zinc metabolism related genes including *MT1E*, *GCLM*, and *GSR* were all down regulated in IPF AEC2s relative to that of AEC2s from healthy lungs (Figure 2A). These data suggested that there was a defect of zinc metabolism with AEC2s from IPF lungs and down regulation of sirtuin signaling of IPF AEC2s might result from zinc metabolic dysfunction of the cells.

**Figure 2.**
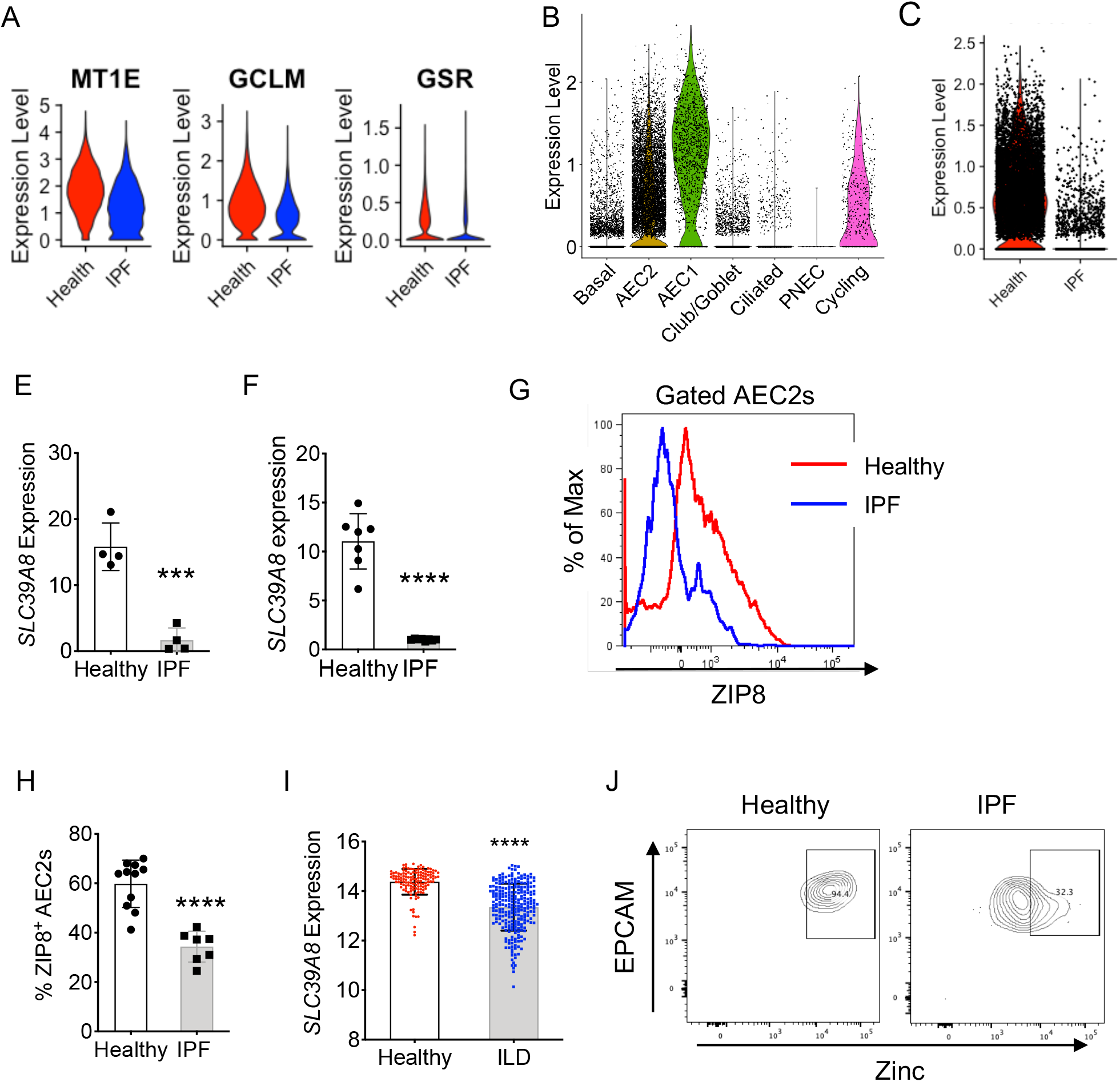
Dysregulation of zinc metabolism of IPF AEC2s. (A) scRNA-seq own regulated zinc metabolism related genes in IPF AEC2s. (B) *SLC39A8* expression of human lung epithelial cell types. (C) *SLC39A8* expression of AEC2s from healthy and IPF lungs. (E) *SLC39A8* expression of AEC2s freshly isolated from lung tissues (n = 4 each. ***p < 0.001). (E) *SLC39A8* expression of AEC2s derived from 3D cultured organoids (n = 7 – 8. ****p < 0.0001). (F, G) Flow cytometry of cell surface ZIP8 level and percent of ZIP8_+_ cells of healthy and IPF AEC2s. (n = 7 – 11, ****p < 0.0001). (H) *SLC39A8* expression of healthy and ILD lung tissues (n = 137 – 255, ****p < 0.0001). (I) Representative plots of intracellular zinc staining of human AEC2s.

Zinc homeostasis is controlled by cell membrane bound zinc transporters (Kambe et al., 2015). Most zinc transporters are tissue specific and cell type specific (Kambe et al., 2015). SLC39 gene family encodes zinc influx transporters, ZIP proteins. There are total 14 members in the ZIP protein family. We looked at the gene expression of ZIP proteins in lung tissue with online deposited datasets (Fagerberg et al., 2014) and found *SLC39A8* (encoding ZIP8) was highly expressed in the lung (Figure S3A). Lung tissue expresses SLC39A8 the highest compared to that of other organs or tissues (Figure S 3B).

Next we tried to determine cell types that express *SLC39A8* in the lung by analyzing our human lung scRNA-seq data. Within lung epithelial cells, *SLC39A8* mainly expressed in alveolar type I cells (AEC1s) and AEC2s (Figure 2B). We then compared *SLC39A8* expression level of AEC2s between healthy and IPF and found AEC2s from IPF lungs expressed much lower *SLC39A8* than that of AEC2s from healthy lungs (Figure 2C). We confirmed decreased *SLC39A8* expression of IPF AEC2s both directly isolated from lung tissues (Figure 2D) and derived from 3D cultured organoids (Figure 2E) using qPCR. We further showed that cell surface expression of ZIP8 protein of IPF AEC2s was decreased compared to that of healthy AEC2s using flow cytometry (Figure 2G, 2H). We analyzed LGRC deposited data and found *SLC39A8* expression was significantly lower in lung tissues of patients with interstitial lung diseases (ILD) (Figure 2I), which is consistent with our findings of decreased *SLC39A8* expression in IPF AEC2s.

*SLC39A8* expression was also decreased in AEC1s from IPF lungs relative to that of AEC1s from healthy lungs (Figure S3C). The significance of ZIP8 in AEC1s is under investigation and not the focus of this study. The expression of *SLC39A1* and *SLC39A7* are relatively high in addition to *SLC39A8* in the lung. We found that the expression of *SLC39A1* was slightly lower in IPF AEC2s (Figure S3D) and *SLC39A7* expression was similar between AEC2s from healthy and IPF lungs (Figure S3D). ZNT family genes were expressed in human AEC2s at a lower level than that of ZIP family genes (Figure S3E). The expression of *SLC30A1, SLC30A6, SLC30A7* are slightly lower in IPF AEC2s compared to healthy AEC2s (Figure S3E). *SLC30A5* and *SLC30A9* expression showed no significant difference between IPF and healthy AEC2s (Figure S3E). Other ZNT family members were not expressed.

Furthermore, IPF AEC2s showed lower intracellular zinc levels compared to that of AEC2s from healthy lungs (Figure 2J). Lower levels of intracellular zinc with IPF AEC2s possibly result from decreased ZIP8 expression of the cells. These data identified a dysregulated zinc matabolism in IPF AEC2s characterized by the loss of *SLC39A8*/ZIP8 expression and lower intracelluar zinc levels.

### ZIP8-dependent zinc metabolism is required for AEC2 renewal

Zinc is an essential element for cell metabolism and involves in various cell functions. It has been reported that zinc metabolism has a role in maintaining epithelial progenitor function (Ohashi et al., 2019; Ohashi et al., 2016; Yang et al., 2020). We hypothesized that ZIP8, as the zinc influx transporter of AEC2s, would play an important role in regulating AEC2 progenitor function. To test this hypothesis, we flow sorted ZIP8_pos_ and ZIP8_neg_ AEC2s from both healthy and IPF lungs, applied the cells to 3D organoid culture, and determined the renewal capacity of AEC2s measured by colony forming efficiency (CFE) (Figure 3A, 3B). ZIP8_neg_ AEC2s showed significantly lower renewal potential compared to that of ZIP8_pos_ AEC2s from the same lung (Figure 3B). Both cells from healthy lungs and IPF lungs showed the same pattern even though ZIP8_pos_ AEC2s from IPF lungs still had lower CFE than the ZIP8_pos_ AEC2s from healthy lungs did (Figure 3B), indicating there might be other factors contribute to impaired renewal of IPF AEC2s. AEC2s derived from colonies of ZIP8_neg_ AEC2s showed reduced *SIRT1* expression relative to that of ZIP8_pos_ AEC2s (Figure 3C) suggesting that zinc homeostasis directly affected sirtuin signaling of AEC2s. ZIP8_neg_ AEC2s also had lower *PDPN* expression indicating that ZIP8 deficiency might also affect AEC2 to AEC1 differentiation (Figure 3D).

**Figure 3.**
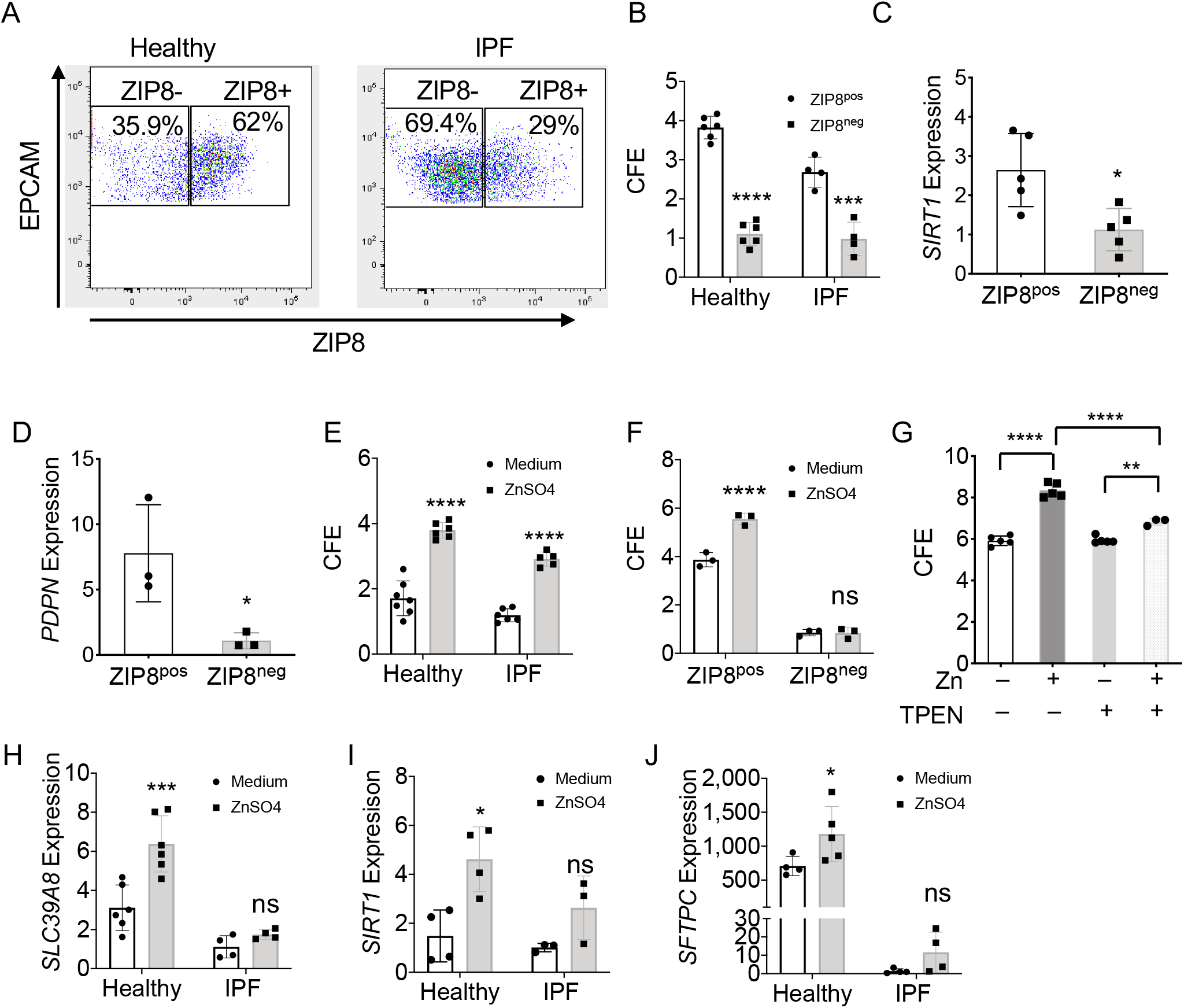
ZIP8 expression is associated with AEC2 renewal. (A) Flow cytometry plots of gated ZIP8_+_ and ZIP8_−_ AEC2s. (B) CFE of flow sorted ZIP8_+_ and ZIP8_−_ AEC2s from healthy and IPF lungs (n = 4 – 6, ***p < 0.001, ****p < 0.0001). (C, D) *SIRT1* (C, n = 5 each, *p < 0.05).) *and PDPN* (D, n = 3 each, *p < 0.05).) expression of AEC2s derived from 3D cultured organoids of ZIP8_+_ and ZIP8_−_ AEC2s from healthy lung. (E) CFE of AEC2s from healthy and IPF lungs with and without zinc sulfate (100 μM) treatment (n = 5 – 7, ****p < 0.0001). (F) CFE of ZIP8_+_ and ZIP8_−_ AEC2s with and without zinc sulfate (100 μM) treatment (n = 3 each. ****p < 0.0001; ns, not significant). (G) CFE of AEC2s with and without zinc sulfate and TPEN treatment (n = 3 – 5, **p < 0.01, ****p < 0.0001). (H-J) Expression of *SLC39A8* (n= 4 – 6, ***p < 0.001), *SIRT1* (n = 3 – 4, *p < 0.05) and *SFTPC* (n = 4 – 5, *p < 0.05) in AEC2s with and without zinc sulfate by RT-PCR (ns, not significant).

Next we tested weather exogenous zinc affects AEC2 progenitor renewal. As proof-of-principle, we applied 100 μM zinc sulfate (ZnSO_4_) into 3D organoid culture of AEC2s and zinc treatment was able to improve the colony forming capacities of AEC2s from both healthy and IPF lungs (Figure 3E). However, zinc treatment showed no effect on CFE of ZIP8_neg_ AEC2s (Figure 3F), indicating ZIP8 might act as a sole functional zinc transport of AEC2s.

To verify the specificity of exogenous zinc treatment on promoting AEC2 progenitor renewal, we applied the zinc chelator, TPEN to the 3D organoid culture. TPEN was able to abolish the effect of zinc treatment in promoting AEC2 renewal (Figure 3G).

Zinc treatment elevated *SLC39A8* expression of AEC2 from both healthy and IPF lungs (Figure 3H). AEC2s derived from 3D organoids with zinc treatment also showed increased *SIRT1* (Figure 3I) and *SFTPC* (Figure 3J) expression. AEC2s from healthy lungs showed much better response to zinc treatment than the AEC2s from IPF lungs did (Figure 3H – 3J).

We have showed previously that a loss of cells surface hyaluronan (HA) of AEC2s in IPF lungs contributed to the impaired renewal capacity of the cells (Liang et al., 2016). Since loss of HA and loss of ZIP8 both happened with IPF AEC2s, we would expect a correlation between ZIP8 expression and cell surface HA levels of AEC2s. We compared HA expression of ZIP8_+_ and ZIP8_−_ AEC2s from both healthy and IPF lungs using flow cytometry and ZIP8_+_ AEC2s showed higher HA levels relative to the ZIP8_−_ AEC2s from the same lung (Figure S4A). Zinc treatment was able to elevate cell surface HA of AEC2s (Figure S4B).

### Reduced renewal capacity of AEC2s from aged mice

Aging is one of the important risk factors to IPF pathogenesis. Next we investigated how aging affect AEC2 progenitor functions. We accessed epithelial cells from the lungs of 2 – 3 mouths old (young) and 18 – 20 months old (aged) mice by flow cytometry using cell surface marker gating strategy established in the lab (Liang et al., 2016). Epithelial cells are CD31_−_CD34_−_CD45_−_EpCAM_+_ (R1), whereas AEC2s are CD31_−_CD34_−_CD45_−_EpCAM_+_ AEC2s (R2) (Figure 4A). The numbers of total cells recovered from each lung were similar between young and old mice (Figure 4B). However, we observed reduced percentage and total number of epithelial cells in old mouse lungs compared to that of young mouse lungs (Figure 4C, 4D). We then looked AEC2s within epithelial population and found the percentage of AEC2s was much lower with the cells from old mouse lungs than the cells from young mouse lungs (Figure 4E), that resulted in extremely low number of AEC2s in the old mouse lungs (Figure 4F).

**Figure 4.**
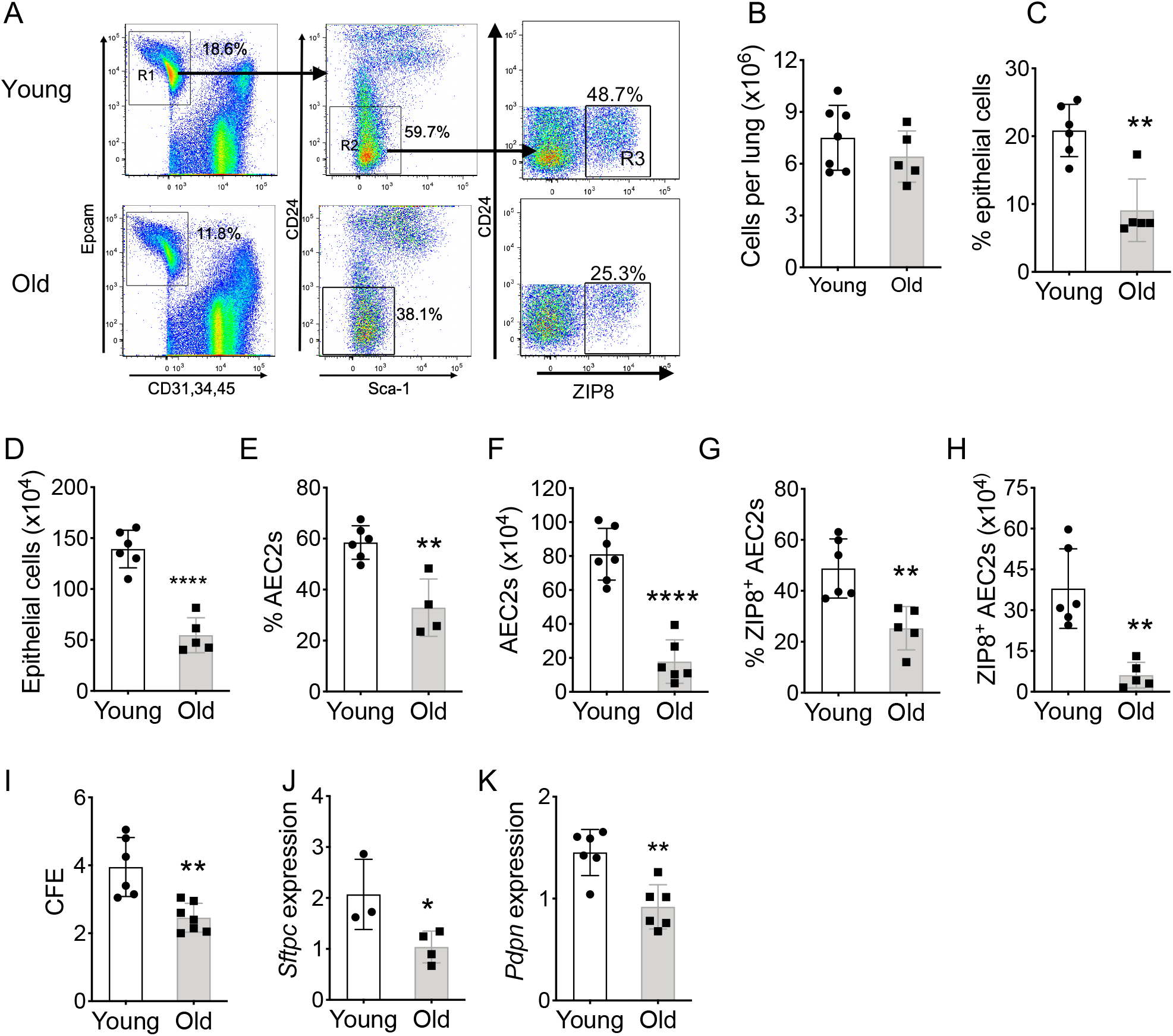
Decreased ZIP8 expression and renewal capacity of aging AEC2s. (A) Gating strategy for ZIP8 expressing AEC2s from young and old mice by flow cytometry. (B) Number of total cells recovered from young and old mouse lungs (n = 5 – 6). (C, D) Percent and number of EpCAM_+_CD31_−_CD34_−_CD45_−_ epithelial cells (R1 in panel A) (n = 5-6. **p< 0.01, ****p < 0.0001). (E) Percent of CD24_−_Sca-1_−_ AEC2s (R2 in panel A) within total lung epithelial cells (R1 in panel A) (n = 4 – 6, **p < 0.01). (F) Number of AEC2s were recovered from young and old mouse lung (n = 6 – 7, ****p < 0.0001). (G) Percent of ZIP8_+_ cells within total AEC2s (R3 in panel A) (n = 5 – 6, **p< 0.01). (H) Number of ZIP8_+_ cells were recovered from young and old mouse lung (n = 5 – 6, **p < 0.01). (I) CEF of mouse AEC2s isolated from young and old mouse lungs (n = 6 – 7, **p < 0.01). (J, K) Expression of *Sftpc* (J) (n = 3 – 4, *p < 0.05) and *Pdpn* (K) (n = 6 each, **p< 0.01) of mouse AEC2s derived from 3D cultured organoids by RT-PCR.

We further analyzed ZIP8 expression of AEC2s and observed a big reduction of percentage (Figure 4A, 4G) and total number (Figure 4H) of ZIP8_+_ AEC2s (R3 in Figure 4A) in old mouse lungs relative to that in young mouse lungs. These data suggested that the loss of ZIP8_+_ AEC2 progenitor cells might be one of the characteristics of lung aging.

Next we compared the regenerative capacity of AEC2s from young and old mouse lungs with 3D organoid culture. AEC2s from old mouse lungs gave rise to fewer colonies than the AEC2s from young mouse lungs (Figure 4I). AEC2s derived from colonies of old mouse AEC2s showed lower expression of *Sftpc* (Figure 4J) and *Pdpn* (Figure 4K) relative to the cells derived from colonies of young AEC2s, indicating impaired renewal and differentiation of AEC2 progenitors with aging.

### Impaired ZIP8-dependent AEC2 renewal in aging and post injury

We have performed single cell RNA-seq of flow enriched lung epithelial cells from both young and old mice at uninjured stage (day 0) as well as day 4 and day 14 after bleomycin injury, which is reported in a separate paper (bioRxiv doi will be updated here). To further investigate the role of aging in regulating ZIP8 expression and progenitor function of AEC2s, we analyzed gene expression of mouse AEC2s from the dataset of mouse lung single cell RNA-seq.

We first performed pathway analysis to compare gene expression program of AEC2s from day 4 bleomycin injured young and old mouse lungs. Similar to what we have observed with IPF AEC2s, we found that the sirtuin signaling pathway was on the top of down regulated pathways with the lowest z-score of AEC2s from old mice relative to AEC2s from young mice (Figure 5A). This finding indicated that injured AEC2s in the injured old mouse lungs might share the similar genomic changes with IPF AEC2s.

**Figure 5.**
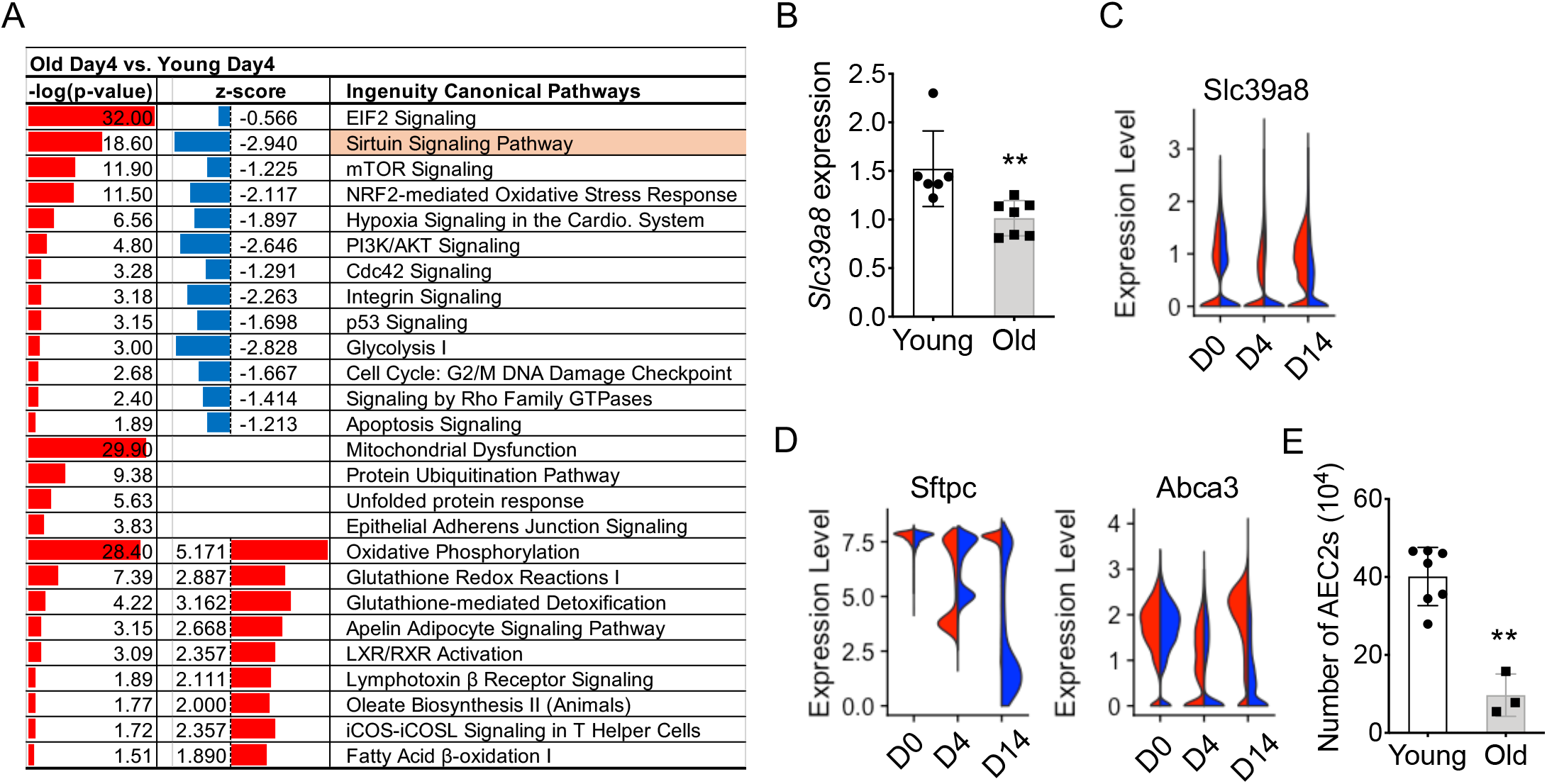
Decreased ZIP8 expression and renewal capacity of aging AEC2s. (A) IPA pathway analysis of AEC2s from young and aged mice 4 days after bleomycin. (B) qPCR *Slc39a8* expression of AEC2s from uninjured mouse lungs (n = 6 – 7, **p< 0.01). (C, D) Expression of *Slc39a8* (C) and AEC2 marker genes *Sftpc* and *Abca3* (D) of AEC2s from day 0 and bleomycin day 4, day 14 young and old mouse lungs. (E) Number of AEC2s were recovered from mouse lung tissue at day 14 after bleomycin injury (n = 3 – 6, ***p< 0.001).

Similar to what we observed with human lung epithelial cells, *Slc39a8* mainly expressed in mouse alveolar epithelial cells (Figure S3F). *Slc39a8* expression was down regulated in AEC2s from old mice relative to that of AEC2s from young mice using q-PCR (Figure 5B).

Next, we analyzed *Slc39a8* expression of AEC2s in the bleomycin injured lungs from young and old mice. At day 4, the timepoint with a maximum of AEC2 injury after bleomycin treatment, *Slc39a8* expression was decreased in AEC2s from both young and old mice (Figure 5C). However, the AEC2s from old mouse lungs had more severe loss of *Slc39a8* expression relative to that of young AEC2s (Figure 5C). It is interesting that at day 14, the epithelium recovery stage after bleomycin injury, the *Slc39a8* expression of ACE2s from young mice was partially recovered, but its expression in AEC2s of old mice remained low (Figure 5C).

Consistent with the pattern of *Slc39a8* expression of AEC2s after bleomycin injury, AEC2 marker genes including *Sftpc* and *Abca3* were highly expressed in both young and old AEC2s from day 0 intact lungs (Figure 5D) and these genes were down regulated in both young and old AEC2s at day 4 after bleomycin injury (Figure 5D). It is most interesting that at day 14, the expression of AEC2 marker genes of AEC2s from young mice was recovered, but their expression of AEC2s from old mouse lungs was lingered behind (Figure 5D). The number of AEC2s recovered from day 14 old mice was significantly low than the number of AEC2s from day 14 young mice (Figure 5E). These data suggest that severe loss ZIP8 of AEC2s in old mouse lung significantly hampered AEC2 recovery of aged mice after lung injury.

### Blunted response to zinc treatment of AEC2s from old mouse lungs

Next, we investigated whether exogenous zinc would affect the progenitor function of murine AEC2s. AEC2s with zinc treatment gave rise to an increased number of colonies in 3D organoid culture relative to AEC2s cultured in control medium (Figure 6A). The increased colony formation of AEC2s with zinc treatment could be explained by elevated ZIP8 expression of the cells (Figure 6B,6C).

**Figure 6.**
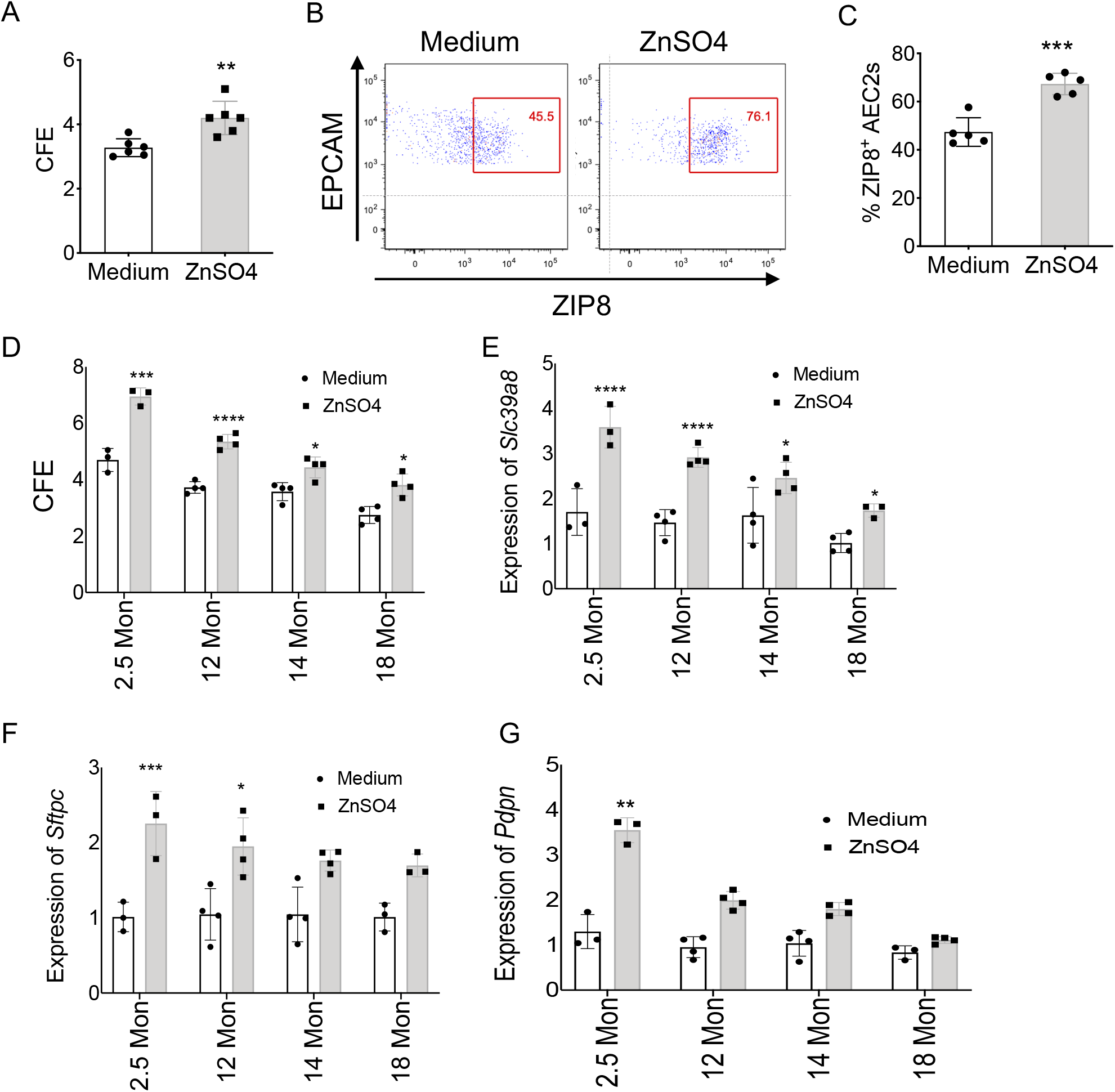
Aging blunted the response of AEC2s to exogenous zinc. (A) CFE of AEC2s from uninjured wild type mice treated with exogenous ZnSO4 (100 μM) (n = 6, **p < 0.01). (B) Flow cytometry gated ZIP8_+_ AEC2s cultured with medium only or medium containing 100 μM ZnSO4. (C) Percent of ZIP8_+_ AEC2s (n = 6, ***p < 0.001). (D) CFE of mouse AEC2s isolated from lungs of 2.5, 12, 14, and 18-month-old mice (n = 3 – 4, *p < 0.05, **p < 0.01, ***p < 0.001, ****p < 0.0001). (E-G) qPCR gene expression of *Slc39a8* (D), *Sftpc* (E), and *Pdpn* (F) (n = 3 – 4, *p < 0.05, **p < 0.01, ***p < 0.001, ****p < 0.0001).

We anticipated that aging would have impact on the responses of murine AEC2s to exogenous zinc treatment. To test this hypothesis, we isolated AEC2s from multiple age groups of mice including 2.5 months old (young) mice, 18 months old (aged) mice, and 12 months and 14 months old in between (Figure 6D – 6G). We applied the flow sorted AEC2s to 3D organoid culture with zinc treatment. Zinc treatment increased the CFEs of AEC2s from all age groups (Figure 6D). However, ACE2s from 2.5 months old young mouse responded to zinc treatment the best and the responses diminished when the mice got older (Figure 6D). *Slc39a8* expression increased in the same pattern as CFE increased with zinc treatment (Figure 6E). AEC2s derived from the organoids with zinc treatment showed increased *Sftpc* (Figure 6F) and *Pdpn* (Figure 6G) expression indicating improved progenitor renewal and differentiation of the cells. The response to zinc treatment was most significant with AEC2s from 2.5 months old (young) mice and the response blunted when the age of the mice increased (Figure 6D – 6G).

### Targeted deletion of Slc39a8 in AEC2s decreased AEC2 renewal

To further illustrate the role of ZIP8 in AEC2 renewal and lung fibrosis in vivo, we generated a mouse line with targeted deletion of *Slc39a8* in the AEC2 compartment, *SFTPC-CreER;Rosa26-tdTomato;Zip8*_*flox/flox*_ (Zip8_ΔAEC2_), by cross breeding *SFTPC-CreER;Rosa26-tdTomato* mice with *Slc39a8* floxed mice (acquired from MMRRC Repository). Rosa26-tdTomato is used to facility cell sorting. Upon tamoxifen treatment, *Slc39a8* expression in Sftpc-expressing cells is deleted. We confirmed the genotype of the of Zip8_ΔAEC2_ mice with PCR (Figure S6A). The Zip8_ΔAEC2_ mice developed normally without obvious physical defect. There are no significant changes in lung histology between 12 weeks old adult Zip8_ΔAEC2_ mice and littermates (Figure S6B).

We assessed AEC2s in the lungs of Zip8_ΔAEC2_ mice and littermate control two weeks after 4 doses of tamoxifen of 200 mg/kg. The ZIP8_+_/Tomato_+_ cells were significantly reduced in the lung of Zip8_ΔAEC2_ mice (Figure 7A, 7C). AEC2s with Zip8 deletion showed decreased intracellular zinc levels compared to that of AEC2s from control mice (Figure 7B). The total number of AEC2s recovered from uninjured lungs was not significantly different between Zip8_ΔAEC2_ mice and littermate controls (Figure 7D). However, AEC2s isolated from uninjured Zip8_ΔAEC2_ mice had reduced *Sirt1* expression (Figure 7E) and lower CFEs with 3D organoid culture than that of AEC2s from littermate controls (Figure 7F).

**Figure 7.**
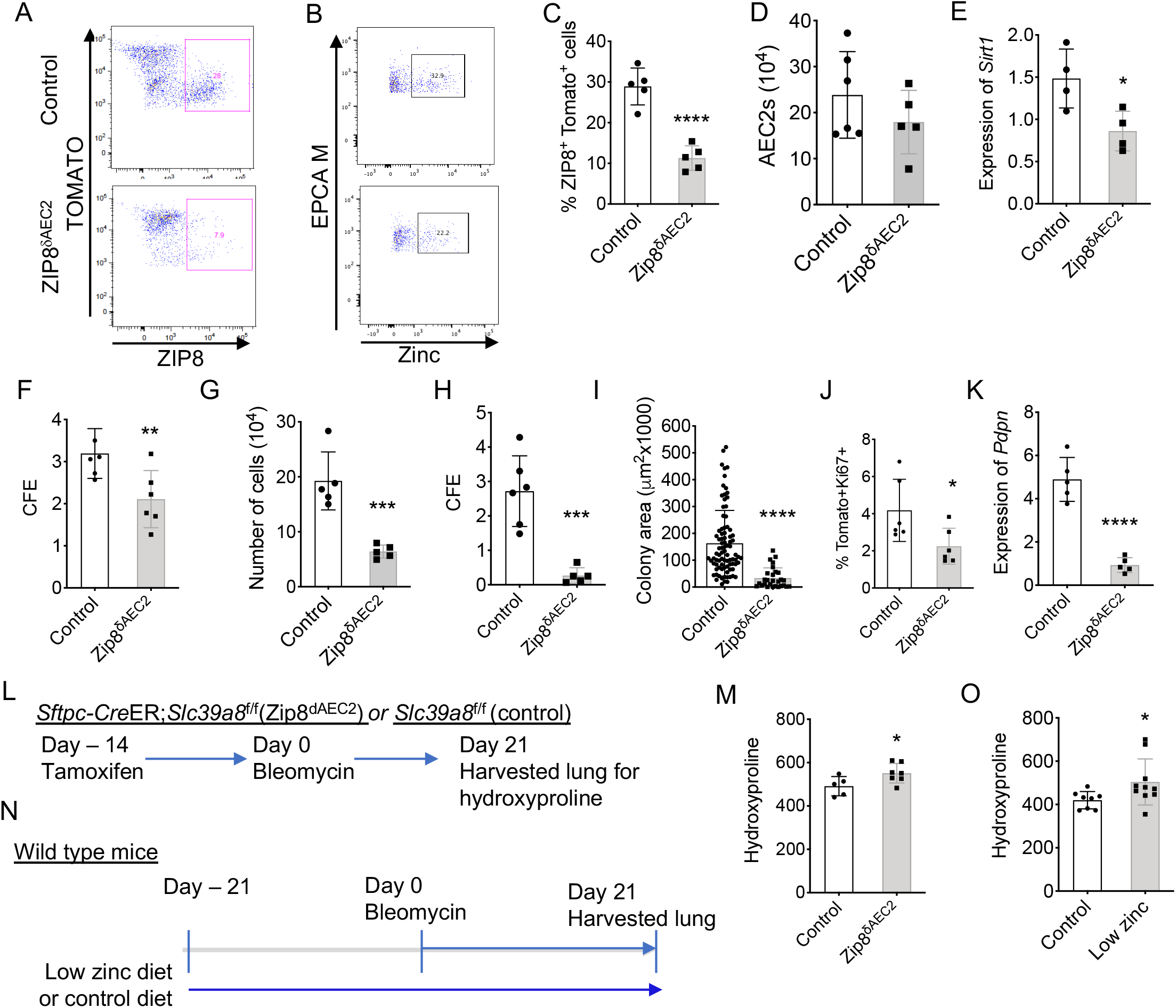
Targeted deletion of *Slc39a8* decreased AEC2 renewal and worsened lung fibrosis. (A) ZIP8 expressing of AEC2s from Zip8_ΔAEC2_ and control mice by flow cytometry. (B) Intracellular zinc in AEC2s from Zip8_ΔAEC2_ (mean 25.03%, n = 4) and control mice (mean 36.10%, n = 6) by flow cytometry (p < 0.01). (C) ZIP8 expressing AEC2s (Tomato_+_) from Zip8_ΔAEC2_ and control mice by flow cytometry (n = 5 each, ****p < 0.0001). (D) Number of AEC2s recovered from uninjured lungs of Zip8_ΔAEC2_ and control mice (n = 6 – 8). (E) *Sirt1* expression of AEC2s by qPCR (n = 4, *p< 0.05). (F) CFE of AEC2s from uninjured mouse lungs (n = 6 each, **p< 0.01). (G) Number of AEC2 recovered per lung of Zip8_ΔAEC2_ and control mice at day 4 after bleomycin (n = 5 each, **p < 0.01). (H, I) CFE (n = 5 – 6) and colony size (n = 28 – 86) of AEC2s isolated from Zip8_ΔAEC2_ and control mice at day 4 after bleomycin (***p< 0.001, ****p< 0.0001). (J) *Ki67* expression of AEC2s derived from 3D cultured organoids of AEC2s from bleomycin injured Zip8_ΔAEC2_ and control mice by flow cytometry (n = 6 each, **p< 0.05). (K) *Pdpn* expression of AEC2s derived from 3D cultured organoids of AEC2s from bleomycin injured Zip8_ΔAEC2_ and control mice by qPCR (n = 5 each, ****p< 0.0001). (L) Experiment layout of Zip8_ΔAEC2_ and control mice treated with 1.25U/kg bleomycin following tamoxifen injection. (M) Hydroxyproline levels of lungs from Zip8_ΔAEC2_ and control mice harvested at day 21 after bleomycin (n = 5 – 7, *p< 0.05). (N) Experiment layout of wild-type mice treated with 2.5 U/kg bleomycin following zinc deficient and control diet. (O) Hydroxyproline levels of lungs of mice fed with zinc deficient diet and control diet harvested at day 21 after bleomycin (n = 8 – 10, *p< 0.05).

Next we treated 8-10 weeks old young adult Zip8_ΔAEC2_ mice and littermate controls with 2.5 U/kg bleomycin following 4 doses of tamoxifen injection and sacrificed the mice at day 4 after bleomycin treatment. The AEC2s recovered from each lung of bleomycin injured Zip8_ΔAEC2_ mice was much fewer than the cells from littermate controls (Figure 7G).

AEC2s from bleomycin injured Zip8_ΔAEC2_ mouse lungs showed severe regenerative defect with 3D organoid culture and they gave rise to extremely fewer and smaller colonies compared to AEC2s from littermate controls (Figure 7H,7I). AEC2s derived from 3D organoids of AEC2s from Zip8_ΔAEC2_ mice showed reduced percentage of Ki67+Tomato+ cells (Figure 7J) and dramatic decrease of *Pdpn* expression (Figure 7K).

### Zinc metabolism regulated lung fibrosis

Next we investigated the role of zinc metabolism in regulating lung fibrosis. We treated Zip8_δAEC2_ mice and control mice with 2.5 U/kg bleomycin after tamoxifen injection and harvested the lungs 21 days after bleomycin injury for fibrosis assessment using hydroxyproline assay. Lungs of bleomycin injured Zip8_δAEC2_ mice showed increased hydroxyproline contents compared to that of the lungs from control mice (Figure 7L, 7M). To further confirm that zinc deficiency worsened lung fibrosis, we applied bleomycin lung injury model to the mice fed with modified zinc diet. We fed the wild type mice with zinc deficient diet (zinc 20 PPM) and control diet (zinc 80 PPM) for three weeks before bleomycin injury (2.5 U/kg) and the mice were kept on the same diet until end of the experiments. We harvested lung tissues at day 21 after bleomycin treatment for fibrosis assessment. Mice fed with zinc deficient diet developed worse lung fibrosis showing as increased hydroxyproline contents compared to the lungs of mice fed with control diet (Figure 7N, 7O). These findings demonstrate that zinc transporter ZIP8 and zinc metabolism have a key role in regulating alveolar progenitor renewal. Reduced zinc transporter ZIP8 of AEC2s in aging and in IPF contributes to impaired alveolar repair and lead to lung fibrosis.

## Discussion

Aging have been recognized as an important risk factor for IPF. Evidences indicates that the main hallmarks of aging occur prematurely in IPF and primarily affect epithelial cells (Rojas et al., 2015; Selman and Pardo, 2014; Thannickal, 2013). However, the molecular mechanism that regulates AEC2 progenitor cell renewal and lung fibrosis during aging and in IPF is poorly understood. In this study we identified reduced expression of *SLC39A8*, the gene encoding zinc transporter ZIP8 in AEC2s from lungs of patients with IPF using single cell transcriptome analysis. We further observed reduced level of cell surface ZIP8 expression and decreased intracellular zinc of IPF AEC2s. Most importantly we showed that ZIP8 plays an important role in regulating AEC2 progenitor renewal and ZIP8_neg_ AEC2s had reduced renewal capacity relative to that of ZIP8_pos_ AEC2s. Zinc sulfate treatment was able to promote renewal of AEC2s from both healthy and IPF lungs. This is the first evidence to show that zinc metabolism homeostasis is crucial for maintaining progenitor function of AEC2s and there is a defect of zinc metabolism with IPF AEC2s.

Our findings with AEC2s from human diseased lung were corroborated well by the results from animal model study in scRNA-seq, in vitro culture, as well as in vivo lung fibrosis. With animal model of aging, we found that the ZIP8 expression and renewal capacity of AEC2s from lungs of old mice were reduced relative to that of AEC2s from young mice. Zinc treatment promoted renewal of AEC2s from young mice while the response to zinc diminished with the AEC2s from older mice. These data demonstrate that decreased ZIP8 expression was associated with aging and that contributed to the impaired renewal capacity of AEC2s. Mice with targeted deletion of ZIP8 in AEC2 compartment had reduced AEC2 renewal and developed more severe lung fibrosis further confirmed the role of ZIP8 in regulating AEC2 progenitor function and lung fibrosis.

There are multiple factors that might contribute to the impaired progenitor function of AEC2s during aging including apoptosis (Korfei et al., 2008), ER stress (Lawson et al., 2011), senescence (Minagawa et al., 2011), and mitochondria dysfunction (Bueno et al., 2015). ZIP8 deficiency added another critical player that contributes to the debilitated progenitor function of AEC2s in the aged lung. Zinc deficiency in elderly has been reported (Bogden, 2004; Prasad et al., 1993). Zinc metabolism may have several ways to influence alveolar progenitor activities. One of the obvious mechanisms is through the zinc-dependent sirtuin signaling. We observed down regulation of sirtuin signaling with IPF AEC2s relative to AEC2s from healthy lungs. With 3D Matrigel organoid culture, we showed that sirtuin activator enhanced and sirtuin inhibitor suppressed colony formation of AEC2s. These data indicated that sirtuin signaling is crucial for AEC2 progenitor renewal through a zinc-dependent mechanism. We further showed that ZIP8_neg_ AEC2s had lower sirt1 expression compared to ZIP8_pos_ cells and zinc treatment elevated sirt1 expression as well as renewal capacity of AEC2s. We concluded that ZIP8 and zinc metabolism regulate AEC2 progenitor function through sirtuin signaling.

In addition to the sirtuin signaling, zinc may have a role in alveolar progenitor and in lung repair through other mechanisms such as regulating antioxidative stress (Tan et al., 2018) and expression of glutathiones (Jiang et al., 1998). We observed a downregulation of NRF2-mediated antioxidative stress as well as glutathione redox response in IPF AEC2s. Furthermore, we have reported previously that loss cell surface hyaluronan (HA) and insufficient IL-6 production contributed to the impaired progenitor function of AEC2s in IPF lungs (Liang et al., 2016). We have examined the correlation between ZIP8 expression and HA in AEC2s. We found that ZIP8 expression is positively correlated with HA levels in AEC2s. ZIP8_pos_ AEC2s showed higher HA level than that of ZIP8_neg_ cells. Zinc treatment increased AEC2 cell surface HA along with ZIP8 and SIRT1 expression. Our data suggest that HA and ZIP8 are both important for AEC2 renewal and loss of both molecules with IPF AEC2s might due to loss epithelial integrity of the cells. Further study is needed to investigate the interaction between zinc metabolism and HA production of AEC2s.

How are aging, zinc metabolism, alveolar progenitor, and fibrosis interconnected? We think that the accumulated events in genetic and epigenetic reprogramming during aging may lead to many key changes such as zinc metabolic dysregulation. Many transcription factors (Liu et al., 2013; Prasad, 2008) and epigenetic mediators such as histone deacetylases (HDACs) (Blanquart et al., 2019; Ren et al., 2017; Seto and Yoshida, 2014) are zinc-dependent. For example, ZIP8 is regulated by NFκB (Liu et al., 2013), whereas zinc regulates NFκB activity (Vasto et al., 2007). Both HDACs and histone H1 polypeptides are the substrates of sirtuins (Seto and Yoshida, 2014; Vaquero et al., 2004). Thus, the zinc defect would further fuel the dysregulation of genetic and epigenetic reprogramming. These compounding factors would further deteriorate the alveolar progenitor, leading to the loss of alveolar epithelial integrity and proper repair after injury.

In summary, we have uncovered a novel mechanism that ZIP8-Sirtuin axis regulates AEC2 renewal. Loss of ZIP8 in AEC2s during aging and in IPF results in disrupted sirtuin function, impaired AEC2 progenitor function, and lung fibrosis. Target zinc metabolism may be a new avenue for developing therapeutics for lung fibrosis.

## Experiment procedures

### Study with cells from lung tissues of human subjects

The use of human tissues for research were approved by the Cedars-Sinai Medical Center Institutional Review Board (Pro00032727), and UCLA Institutional Review Board (13-000462-AM-00019). Informed consent was obtained from each subject.

### Mice

All animal experiments were approved by the Institutional Animal Care and Use Committee at Cedars-Sinai Medical Center (IACUC008529). All mice were housed in a pathogen-free facility at Cedars-Sinai Medical Center. *SFTPC-CreER* mice and *Rosa-Tomato*_*flox/flox*_ mice were described previously (Barkauskas et al., 2013; Liang et al., 2016). *Slc39a8*_*flox*/+_ was generated using the Cre/loxP system and acquired from MMRRC Repository. All mice have been backcrossed on C57Bl/6J background more than six generations. Eight to 12 weeks old (young) and 18 months old (aged) wild-type C57Bl/6J mice were obtained from The Jackson Laboratory and housed in the institution facility at least 2 weeks before experiments. Animals were randomly assigned to treatment groups, and they were age- and sex-matched.

### Bleomycin instillation

Bleomycin instillation was described previously (Liang et al., 2016). Under anesthesia, the trachea was surgically exposed. 1.25 – 2.5 U/kg bleomycin (Hospira, Lake Forest, IL) in 25 μl PBS was instilled into the mouse trachea with a 25-G needle inserted between the cartilaginous rings of the trachea. Control animals received saline alone. The tracheostomy site was sutured, and the animals were allowed to recover. Mice were sacrificed at different time points and lung tissue were collected for experiments.

### Hydroxyproline

Collagen contents in mouse lungs were measured with a conventional hydroxyproline method (Jiang et al., 2004; Liang et al., 2019). In brief, lung tissues were vacuum dried and hydrolyzed with 6N hydrochloride acid at 120_o_C for overnight. Hydroxyproline content was measured and expressed as mg per lung. The ability of the assay to completely hydrolyze and recover hydroxyproline from collagen was confirmed using samples containing known amounts of purified collagen.

### Lung tissue and histology

Mice were sacrificed at various time points with or without bleomycin treatment under anesthesia. The trachea was cannulated, and the lung was inflated with 1.0 mL of 10% neutral buffered formalin. Mouse lung tissue was fixed, embedded in paraffin, sectioned to 5 μm slices for H&E and and Masson’s trichrome staining (Geng et al., 2019; Jiang et al., 2004; Liang et al., 2019).

### Cell lines

Mouse lung fibroblast cell line, MLg2908 (Catalog CCL-206), was from ATCC (Manassas, VA). Mycoplasma contamination was assessed with a MycoFluor™ Mycoplasma Detection Kit (Catalog M7006, Thermo Fisher Scientific) and cells used for experiments were free of mycoplasma contamination.

### RNA analysis

RNA was extracted from mouse AEC2s or human AEC2s using TRIzol Reagent. For real-time PCR analysis, 0.5 μg total RNA was used for reverse transcription with the High Capacity cDNA Reverse Transcription Kit (Applied Biosystems, Carlsbad, California). One microliter cDNA was subjected to real-time PCR by using Power SYBR Green PCR Master Mix (Applied Biosystems) and the ABI 7500 fast Real-Time PCR system (Applied Biosystems). The specific primers were designed on the basis of cDNA sequences deposited in the GenBank database: Human *SLC39A8* (NM_001135146) forward CATCTGTCCAGCAGTCTTACAGC, and reverse GACAGGAATCCATATCCCCAAACT. Mouse *Slc39a8* (NM_001135149): forward AGCGATCCTGTGTGAGGAGT and reverse CGGAGAGGAAGTTGAACAGC. Human *SFTPC* (NM_003018.4): forward ATCCCCAGTCTTGAGGCTCT and reverse CTTCCACTGACCCTGCTCAC. Mouse *Sftpc* (NM_011359.2): forward GCAGGTCCCAGGAGCCAGTTC and reverse GGAGCTGGCTTATAG GCCGTCAG. Human *PDPN* (NM_006474.5): forward GGAAGGTGTCAGCTCTGCTC and reverse CGCCTTCCAAACCTGTAGTC. Mouse *Pdpn* (NM_010329.3): forward GCACCTCTGGTACCAACGCAGA and reverse TCTGAGGTTGCTGAGGTGGACAGT. Human *SIRT1* (NM_012238.5): forward TGCCGGAAACAATACCTCCA and reverse AGACACCCCAGCTCCAGTTA. Mouse *Sirt1* (NM_019812.3): forward GAGCTGGGGTTTCTGTCTCC and reverse CTGCAACCTGCTCCAAGGTA. Human *GAPDH* (NM_002046) forward, CCCATGTTCGTCATGGGTGT and reverse, TGGTCATGAGTCCTTCCACGATA. Mouse *Gapdh* (NM_00100130): forward ATCATCTCCGCCCCTTCTG and reverse, GGTCATGAGCCCTTCCACAAC. The relative expression level of each gene was determined against GAPDH level in the same sample. The fold-change of the target genes was calculated by using the 2_−ΔΔCT_ method.

### Mouse lung dissociation and flow cytometry

Mouse lung single cell suspensions were isolated as previously described (Liang et al., 2016). In brief, lungs were perfused with 10 ml PBS and then digested with 4 U/ml elastase (Worthington Biochemical Corporation, NJ) and 100 U/ml DNase I (Sigma), and resuspended in Hanks’ balanced saline solution supplemented with 2% fetal bovine serum (FBS), 10 mM HEPES, 0.1 mM EDTA (HBSS+ buffer). The cell suspension was incubated with primary antibodies including CD24-PE, EpCAM-PE-Cy7, Sca-1-APC, biotinylated-CD31, -CD34, and -CD45, as well as conjugated ZIP8 (rabbit IgG) for 45 minutes. Biotin-conjugated antibodies were detected following incubation with streptavidin-APC-Cy7 (catolog 405208, BioLegend, San Diego, CA), while ZIP8 was detected with goat anti rabbit IgG-FITC for 30 minutes. Dead cells were discriminated by 7-amino-actinomycin D (7-AAD) (BD Biosciences, San Diego, CA) staining. Flow cytometry was performed using a Fortesa flow cytometer and FACSAria III sorter (BD Immunocytometry Systems, San Jose, CA) and analyzed using Flow Jo 9.9.6 software (Tree Star, Ashland, OR).

Primary antibodies EpCAM-PE-Cy7 (clone G8.8, Catalog # 118216, RRID AB_1236471) and Ki-67 (clone 16A8, catalog # 652403, AB_2561524) were from BioLegend. CD24-PE (clone M1/69, Catalog # 12-0242-82, RRID AB_467169), Sca-1 (Ly-6A/E)-APC (clone D7, Catalog # 17-5981-82, RRID AB_469487), CD31 (PECAM-1) (clone 390, Catalog # 13-0311-85, RRID AB_466421), CD34 (clone RAM34, Catalog # 13-0341-85, RRID AB_466425), and CD45 (clone 30-F11, Catalog # 13-0451-85, RRID AB_466447) were all from eBioscience (San Diego, CA). SLC39A8 polyclonal antibody (Catalog # PA5-26368, RRID AB_2543868) was from ThermoFisher.

### Human lung dissociation and flow cytometry

Human lung single cell isolation and flow cytometer analysis were performed as described previously (Barkauskas et al., 2013; Liang et al., 2016). In brief, human lung tissues were minced and then digested with 2 mg/ml dispase II, followed by 10 U/ml elastase and 100 U/ml DNase I digestion. Finally, cells were filtered through 100 μm cell strainer, and lysed with red blood cell lysis to get single cell suspension. Antibody staining was similar as with mouse cells. Flow cytometry was performed with Fortesa and FACSAria III flow cytometer and analyzed with Flow Jo 9.9.6 software. Anti-human CD31 (clone WM59, Catalog # 303118, RRID AB_2247932), CD45 (clone WI30, Catalog # 304016, RRID AB_314404), EpCAM (clone 9C4, Catalog # 324212, RRID AB_756086) were from BioLegend. SLC39A8 polyclonal antibody (Catalog # PA5-26368, RRID AB_2543868) and goat anti-mouse IgG/IgM (Catalog # A-10680, RRID AB_2534062) were from ThermoFisher. HTII-280 (Gonzalez et al., 2010) was a gift from Dr. L. Dobbs lab at UCSF.

### Intracellular staining for zinc of human and mouse AEC2s

Zinc assay kit (catalog # ab241014) was from Abcam (Cambridge, MA). Single cells isolated from human and mouse lungs were cultured on collagen IV coated plated with 100 μm ZnSO4 for overnight. Cells were first stained for intracellular zinc following manufacturer’s protocol, then AEC2 cell surface markers were stained. Flow cytometry was performed using a Fortesa flow cytometer. Intracellular zinc staining of gated human or mouse AEC2s was analyzed using Flow Jo 9.9.6 software (Tree Star, Ashland, OR).

### Matrigel culture of human and mouse AEC2s

Flow sorted human (EpCAM_+_HTII-280_+_) or mouse (EpCAM_+_CD24_−_Sca-1_−_) AEC2s were cultured in Matrigel/medium (1:1) mixture in presence of lung fibroblasts MLg2908 cells (Barkauskas et al., 2013; Chen et al., 2012a; Liang et al., 2016). 100 μl Matrigel/medium mix containing 3 × 10_3_ AEC2s and 2 × 10_5_ MLg2908 cells (Catalog CCL-206, ATCC, Manassas, VA) were plated into each 24 well 0.4 mm Transwell inserts. 400 μl of medium were added in the lower chambers. Cells were cultured with medium alone or with the treatments indicated. Medium was described previously (Liang et al., 2016). Matrigel (growth factor reduced basement membrane matrix) (catalog # 354230) was from Corning Life Sciences (Tewksbury, MA). For cell treatment, the following chemicals were used. 100 μM ZnSO4, 1 mM TPEN, , and 150 mM splitomicin were from Sigma (St. Louis, MO). 1 mM SRT1720 was from Selleck Chemicals (Houston, TX). Same volume of DMSO was used as control. Fresh medium was changed every other day. Cultures were maintained in humidified 37_o_C and 5% CO_2_ incubator. Colonies were visualized with a Zeiss Axiovert40 inverted fluorescent microscope (Carl Zeiss AG, Oberkochen, Germany). Number of colonies with a diameter of ≥ 50 μm from each insert was counted and colony-forming efficiency (CFE) was determined by the number of colonies in each culture as a percentage of input epithelial cells at 12 day after plating (dpp).

### Organoid size measurement

Organoids derived from flow sorted Rosa-Tomato_+_ cells from *SFTPC-CreER*_+_;*Rosa-Tomato*_*flox/flox*_ mice and *SFTPC-CreER*_+_;*Rosa-Tomato*_*flox/flox*_; *Slc39a8*_*flox/flox*_ (Zip8_δAEC2_) mice were pictured at day 12 after plating using Zeiss Axiovert40 inverted fluorescent microscope. ZEN pro 2012 software (Zeiss) was used for measure surface area of the pictured organoids same as previously described (Liang et al., 2016).

### scRNA-seq

scRNA-sequencing was performed in Cedars-Sinai Medical Center Genomic Core. In brief, flow sorted human and single cells were lysed, mRNA was reverse transcribed and amplified as previously described (Xie et al., 2018). Barcoding and library preparation were done with standard procedures according to manufacture manuals (10X Genomics, Pleasanton, CA, USA). The barcoded libraries were sequenced with NextSeq500 (Illumina, San Diego, CA, USA) to obtain a sequencing depth of ~200K reads per cell.

Raw scRNA-seq data was aligned to human genome GRCh38 and mouse genome mm10 with Cell Ranger (10X Genomics) respectively. Downstream quality control, normalization and visualization were done with Seurat package. For quality control, the output expression matrix from Cell Ranger was done based on number of genes detected in each cell, number of transcripts detected in each cell and percentage of mitochondrial genes. The expression matrix was then normalized and visualized with UMAP.

Ingenuity Pathway Analysis (IPA) was performed as described previously (Kramer et al., 2014). Differential expression genes with logFC over 0.1 between healthy and IPF AEC2s were analyzed with R software. The data was then imported into IPA software (Qiagen, Hilden, Germany). Canonical pathways from the core analyze were further analyzed.

### Statistics

The statistical difference between groups in the bioinformatics analysis was calculated using the Wilcoxon Signed-rank test. For the scRNA-seq data the lowest p-value calculated in Seurat was p < 2.2e-10-16. For all other data the statistical difference between groups was calculated using Prism (version 8.4.3) (GraphPad Software, San Diego, CA) and the exact value was shown. Data are expressed as the mean ± SEM. All experiments were repeated two or more times. Data were normally distributed and the variance between groups was not significantly different. Differences in measured variables between experimental and control group were assessed using Student’s t-tests. One-way followed by Bonferroni’s or two-way ANOVA followed Tukey’s multiple comparison test was used for multiple comparisons. Results were considered statistically significant at p < 0.05.

## Acknowledgments

The authors thank the members of our laboratory for support and helpful discussion during the course of the study. This work was supported by National Institutes of Health grants R35-HL150829, R01-HL060539, R01-AI052201, R01-HL077291 (PWN), and R01-HL122068 (DJ and PWN), and P01-HL108793 (PWN and DJ). SCR was supported by European Union’s Horizon 2020 research and innovation programme under the Marie Skłodowska-Curie grant agreement 797209. We thank Dr. L. Dobbs of UCSF providing antibodies for the study.

## Author Contributions

JL, DJ, and PWN conceived the study. JL performed most of the experiments and analyzed the data. GH analyzed single cell RNA transcriptome data, performed flow cytometry analysis, and prepared figures. JL, GH, XL, YW, ND, CY, and DJ analyzed single cell RNA transcriptome data. XL, FT, NL, AB, SR, and TX took part in mouse, cell culture, and biological experiments. SSW and JB provided human samples and interpreted data. BS and WCP interpreted data and contributed with comments on the manuscript. JL, DJ, and PWN wrote the paper. All authors read and reviewed the manuscript.

## Lead Contact Statement

Further information and requests for resources and reagents should be directed to and will be fulfilled by the Lead Contact, Paul Noble.

## Materials Availability

All unique/stable reagents generated in this study are available from the Lead Contact with a completed Materials Transfer Agreement.

## Data and Code Availability

The deposition of the raw data files of the single cell RNA-seq are in progress. The accession numbers for the the RNA-seq analyses and the dataset information will be updated accordingly. R code files used for data integration and analysis are available at https://github.com/jiang-fibrosis-lab.

---

**Figure S1.**
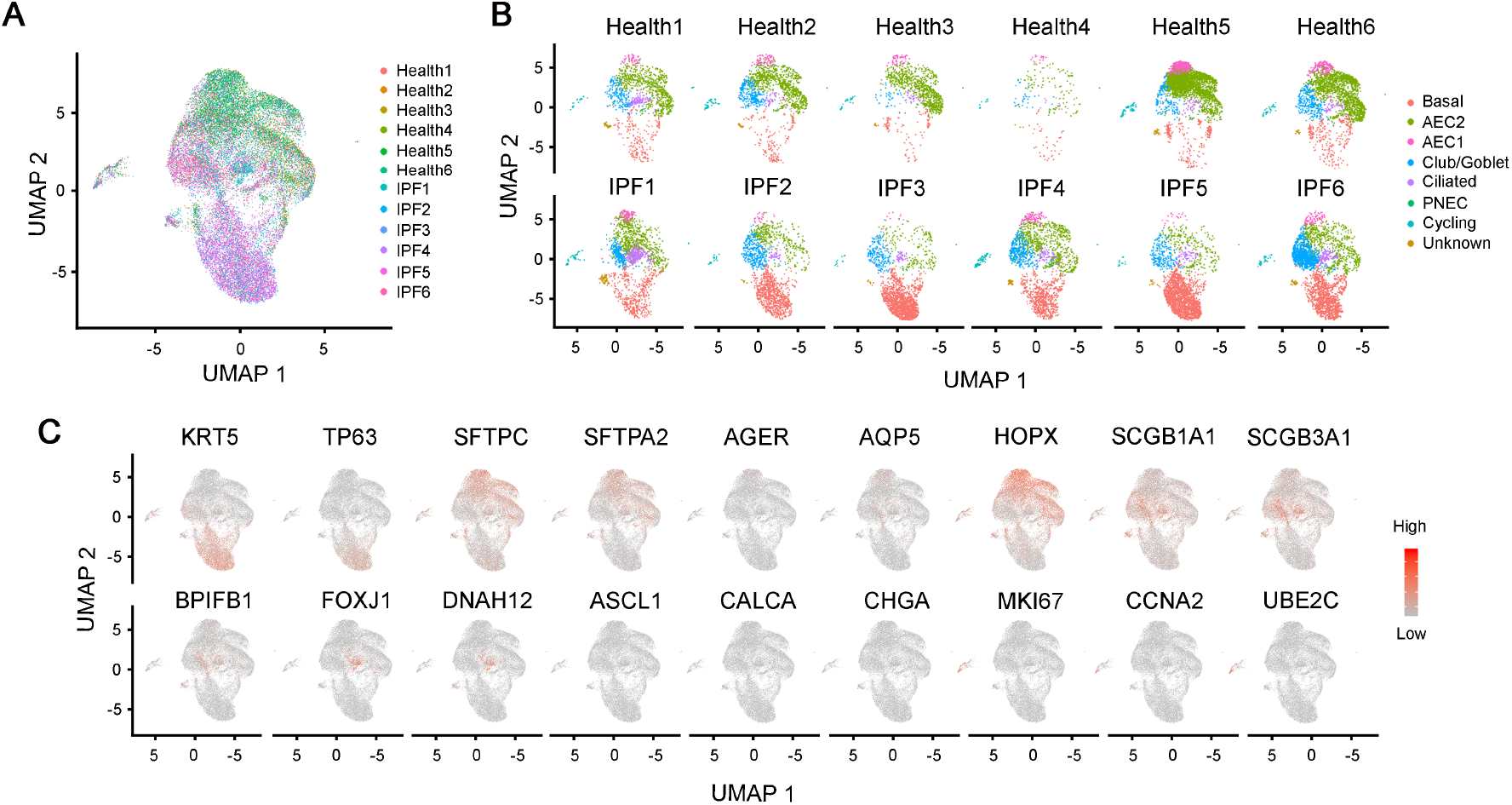
scRNA-seq analysis of lung epithelial cells. (A) UMAP plots of lung epithelial cells from IPF and healthy donors (n = 6 each). (B) Feature plots of lung epithelial clusters grouped by subject. (C) Feature plots of lung epithelial clusters by classic cell type markers.

**Figure S2.**
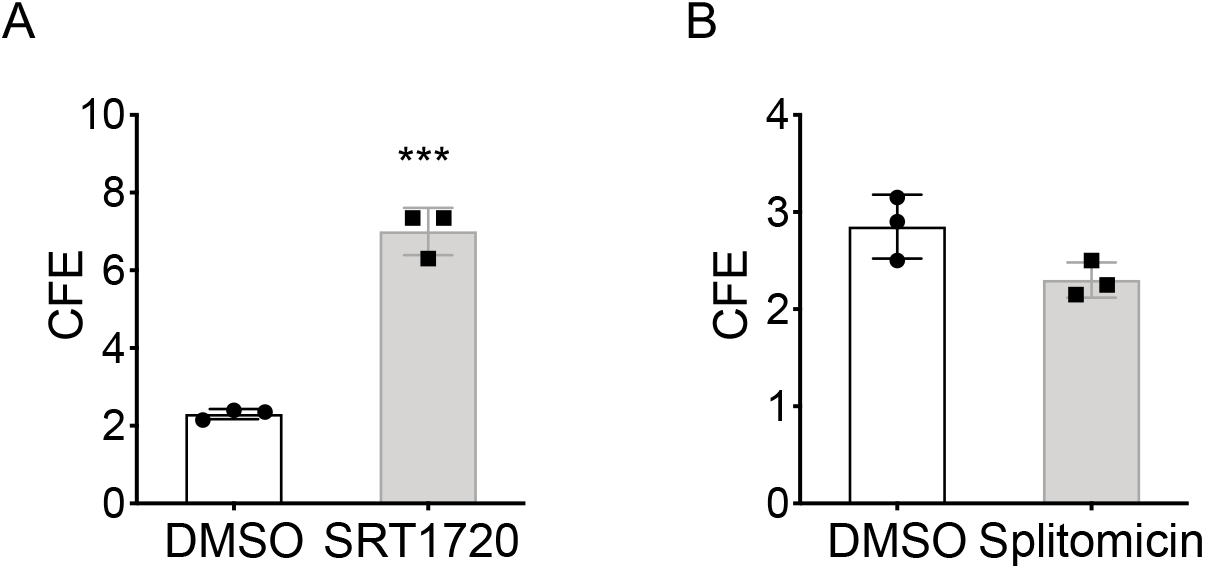
Effect of SIRT activator and inhibitor on mouse AEC2s. (A,B) CFE of AEC2 from uninjured wild type mice treated with SRT 1720 (A), splitomicin (B) with DMSO as control (n = 3 each. ***p< 0.001).

**Figure S3.**
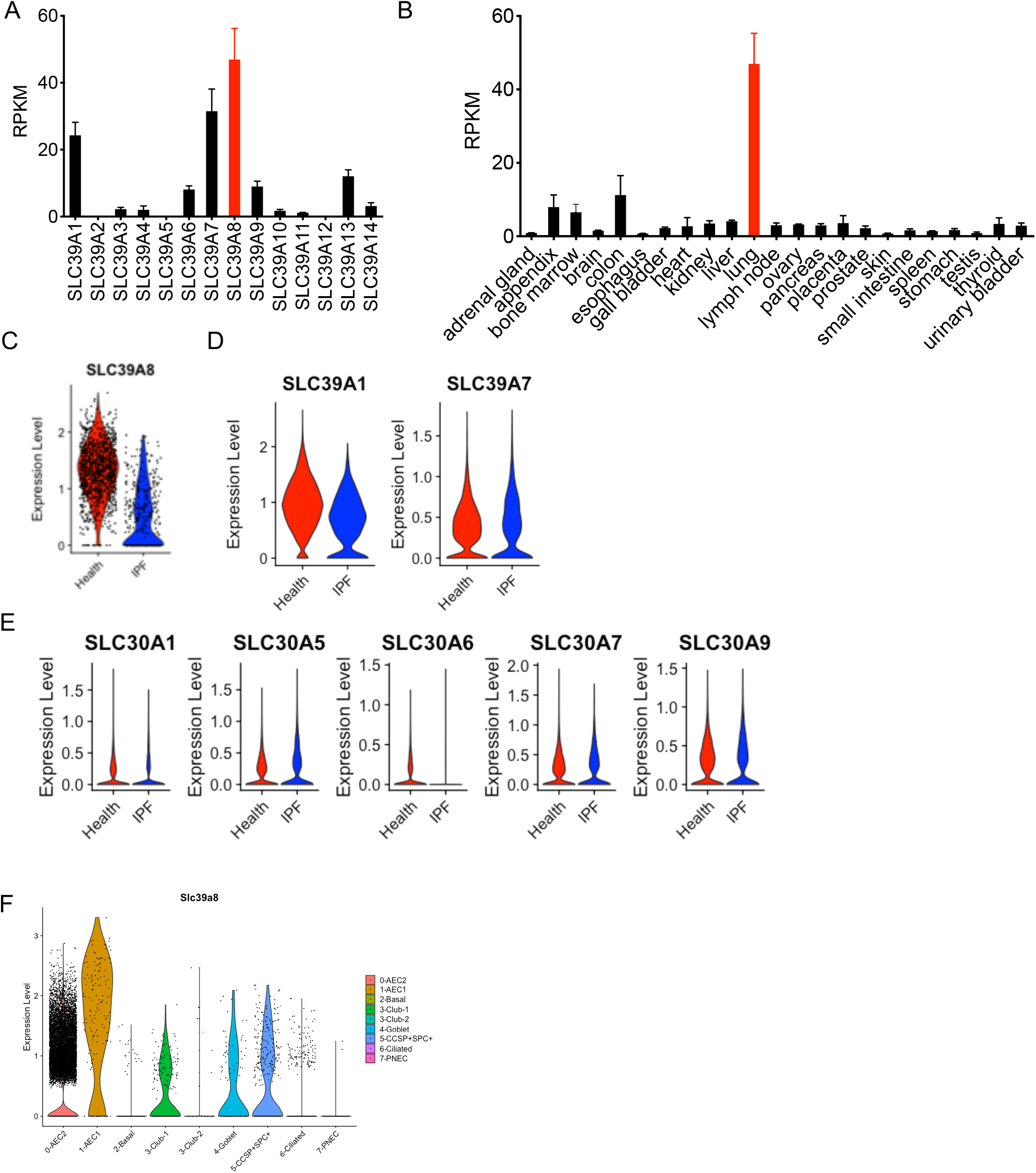
Specific expression of SLC39A8 in alveolar epithelial cells. (A) Expression of *SLC39A1-14* in lung tissue. (B) Expression of *SLC39A8* in human tissues and organs. (C) Expression of *SLC39A8* in AEC1s from healthy and IPF lungs. (D) Expression of *SLC39A1* and *SLC39A7* in AEC2s from healthy and IPF lungs. (E) Expression of ZNT family genes *SLC30A1, SLC30A5, SLC30A6, SLC30A7*, and *SLC30A9* in AEC2s from healthy and IPF lungs. (F) Expression of *Slc39a8* in mouse lung epithelial cells.

**Figure S4.**
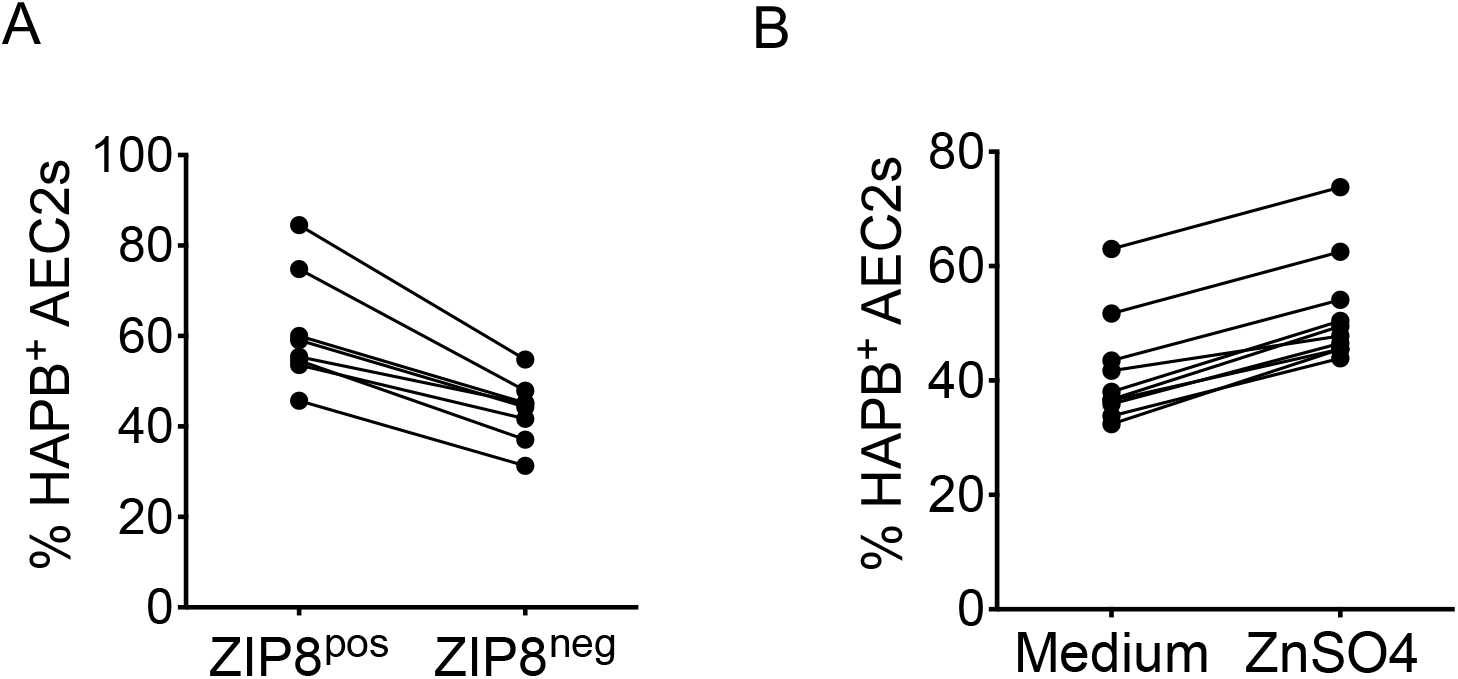
Correlation of ZIP8 and hyaluronan expression. (A) Percent of HABP_+_ of ZIP8_neg_ and ZIP8_pos_ ACE2s by flow cytometry. (B) Percent of HABP_+_ AEC2s with and without 100 μm ZnSO4 treatment.

**Figure S5.**
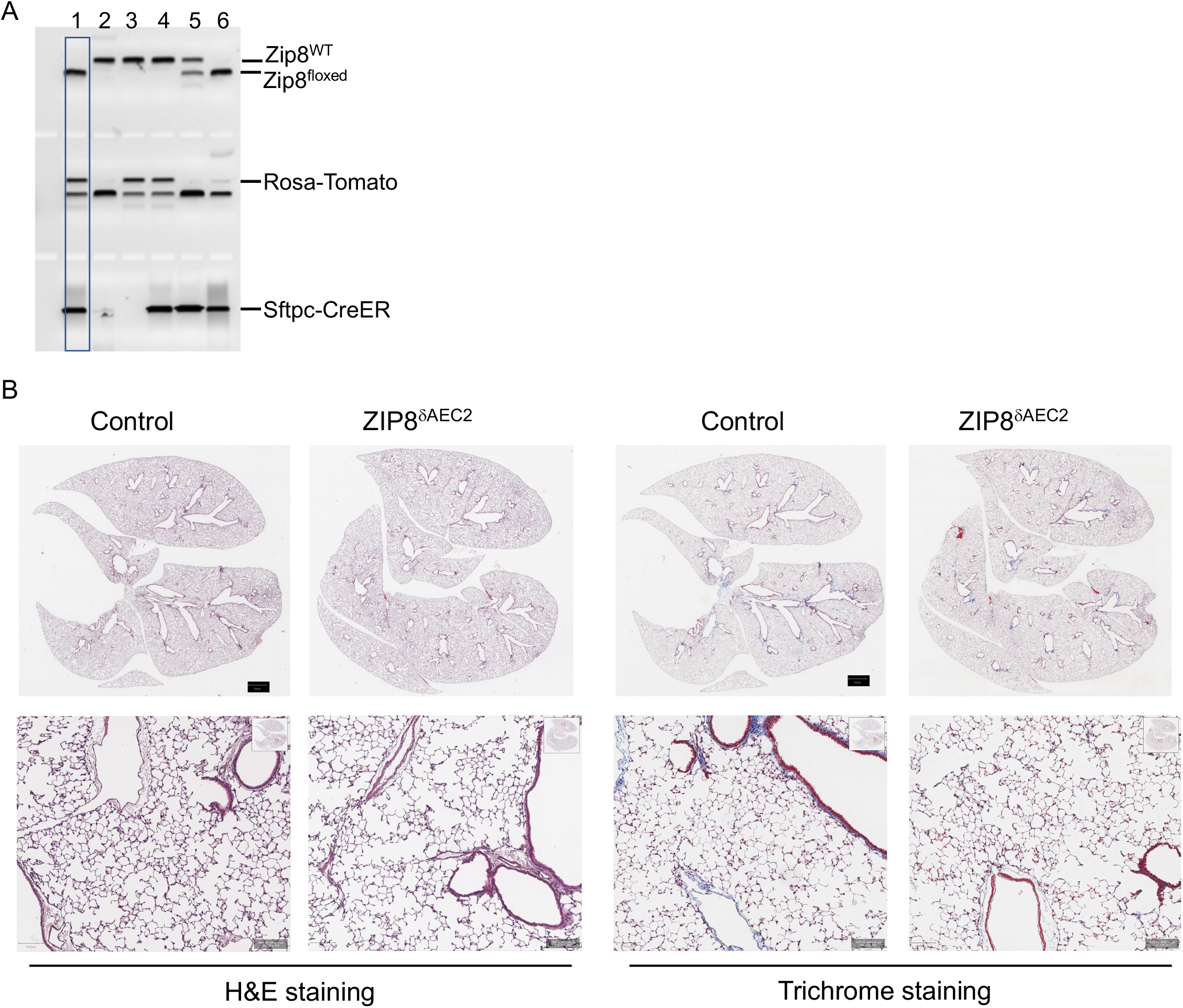
No obvious morphological defect in Zip8 deletion mice. (A) A representative genotyping result of Sftpc-CreER_+_;RosaTomato_+_;Zip8_flox/flox_ (Zip8_ΔAEC2_) mice. (B,C) Histological characterization (B, H&E staining; C, trichrome) of uninjured lungs of Zip8_ΔAEC2_ and control mice at 8 – 10 weeks old.

